# Spatial mapping of rheumatoid arthritis synovial niches reveals specific macrophage networks associated with response to therapy

**DOI:** 10.1101/2023.10.20.563040

**Authors:** Julien De Lima, Marie-Astrid Boutet, Olivier Bortolotti, Laure-Agnès Chépeaux, Yaël Glasson, Anne-Sophie Dumé, Adrien Le Pluart, Alessandra Nerviani, Liliane Fossati-Jimack, Henri-Alexandre Michaud, Jérôme Guicheux, Benoit Le Goff, Costantino Pitzalis, Gabriel Courties, Florence Apparailly, Frederic Blanchard

## Abstract

Rheumatoid arthritis (RA) is a chronic inflammatory autoimmune disease affecting peripheral joints and for which approximately 40% of the patients respond insufficiently to the available synthetic or biologic disease modifying anti-rheumatic drugs (DMARDs). The infiltration of the synovial membrane by lymphocytes and monocytes profoundly alters its homeostatic functions, leading to chronic joint inflammation and bone destruction. A better understanding of how DMARDs impact the complex synovial cell social network in relationship to response/ non-response remains an unmet need to design more targeted and active therapeutic strategies. Here, we used imaging mass cytometry (IMC) to comparatively profile more than 115,000 cells in the synovial tissue of healthy, low inflammatory osteoarthritis and matched active early treatment-naïve RA patients at baseline and at 6 months after starting DMARDs treatment. We notably highlighted that tissue resident macrophages (LYVE1^+^CD206^+^) in perivascular synovial niches encompassing specific subsets of vascular cells, fibroblasts and immune cells vanished in active RA but were recovered in response to DMARDs treatment. Combined ligand-receptor analysis of single-cell RNA sequencing datasets identified that IL10, C-type lectin or TAM (TYRO3, AXL and MERTK) receptors were particularly involved in the restoration of these spatial cell interactions in the context of clinical remission. In addition to providing an unprecedented synovial spatial mapping, our work uncovered novel potential cellular and molecular targets for the development of therapies for RA.

**One Sentence Summary:** Single-cell spatial profiling of rheumatoid arthritis synovium identifies specific cell states linked to treatment response

## INTRODUCTION

Rheumatoid Arthritis (RA) is the most common form of inflammatory arthritis, affecting 0.3-1% of the general population worldwide and characterized by joint inflammation associated with debilitating bone and cartilage erosions. Despite the availability of a large therapeutic arsenal, including conventional synthetic (cs) disease-modifying antirheumatic drugs (DMARDs; e.g., methotrexate, MTX), biologic (b) DMARDs (e.g., TNF inhibitors) and targeted synthetic (ts) DMARDs (e.g., JAK inhibitors), around 40% of patients do not respond adequately to the available treatments, and RA remains a major public health challenge (*1, 2*). RA is a prototypic synovitis-driven autoimmune disease and the in-depth characterization of the synovial tissue have broadened our understanding of RA pathogenesis and led to the development of targeted therapies. Blood biomarkers of inflammation and autoimmunity e.g. ESR / CRP, Rheumatoid Factor / Anti Citrullinated Protein Antibody (ACPA), and clinical features are the mainstay for the diagnosis of RA. However, there are no biomarkers predictive of treatment-response, consequently treatment allocation is based on “trial and error” (*3, 4*). On the other hand, similarly to other rheumatic conditions such as lupus nephritis, vasculitis and myositis, recent evidence in RA supports the concept of integrating synovial cellular/molecular signatures into clinical algorithms to help predicting response to specific DMARDs and enhance clinical outcome, towards precision medicine (*1, 5*).

Also recently, single cell profiling of dissociated tissues using transcriptomic (scRNAseq) and cytometry by time-of-flight (CyTOF) approaches, have brought unprecedented insights into synovial cell heterogeneity, including fibroblasts, macrophages, T and B cells (*6, 7*). Importantly, single cell studies have highlighted the existence of key cellular crosstalk and dysregulated signaling pathways involved in RA pathogenesis. For instance, NOTCH3 signaling appears critical in the differentiation of CD90^+^ perivascular and sublining fibroblasts, and is involved in inflammation and joint damage (*8*). In addition, several T helper cell subsets are also found expanded in the RA synovium, within so called ectopic tertiary lymphoid structures (ELS), to promote B-cell responses (*9*). A particular role for tissue resident macrophages (TRMs) expressing *MERTK, CD163, TREM2* and *TIMD4* (and *Cx3cr1* in mice) have also been proposed as a protective tight-junction-forming cell layer that secludes the synovium, and avoid the infiltration of inflammatory cells from the joint cavity (*7, 10*). Overall, TRMs represent the majority of lining and sublining synovial macrophages in healthy donors and RA patients in remission, whereas several inflammatory macrophage subsets (IMs), characterized by TLR activation, interferon signature, alarmin production or antigen-presenting cell profile, are enriched in the synovial tissue of patients with active RA (*11*).

Although these findings have been outstandingly informative, tissue dissociation represents a major limitation inherent to single cell profiling approaches, as critical cells might be lost in the elution process and spatial information is also lost. This is important, as spatial function/identity, both for the stromal microenvironment and immune cell infiltration are increasingly recognized as key for pathogenesis and treatment response in rheumatology, as in oncology (*5, 12–14*). For example, in oncology, in depth spatial molecular imaging revealed that the proximity of cytotoxic T cells to melanoma cells is associated with positive response to immune checkpoint inhibitors and that specific cellular neighborhoods composed of myeloperoxidase-positive macrophages are associated with long-term survival in patients with glioblastoma (*12, 13*). However, a precise spatial characterization of individual cells within their synovial niche using highly multiplexed histology has neither yet been performed in RA, in comparison with healthy or disease control synovial tissue.

Here, we performed single-cell high dimensional imaging mass cytometry (IMC) to spatially analyze the expression of 33 proteins at subcellular resolution, comparatively in the synovial tissue of healthy individuals, osteoarthritis (OA) patients who lack severe inflammatory responses, and matched samples from active early treatment-naïve RA patients at baseline and at 6 months after starting a DMARD, namely MTX, treatment. For the first time, we provided a comprehensive characterization of synovial cells in the lining, sublining and ELS niches across healthy and disease conditions. We found that LYVE1^+^CD206^+^ TRM population was disturbed in the synovium of early treatment-naive patients and significantly reestablished in a distinct cellular neighborhood following MTX treatment. Furthermore, using ligand-receptor analysis strategies of public scRNAseq databases, we highlighted that LYVE1^+^CD206^+^ TRM cell interactions specifically observed in responder RA patients may involve several mediators, such as IL10, C-type lectin receptors and TAM receptors (TYRO3, AXL and MERTK). Overall, this study provides insights into the spatial distribution and the pathogenic connections between synovial niches and highlights key pathways relevant for better understanding RA pathogenesis and for the development of novel therapeutic strategies.

## RESULTS

### Highplexed imaging mass cytometry highlights high heterogeneity of TRMs within the lining and sublining layers of the synovium

To comprehensively profile the cellular composition and spatial organization of RA synovium, we took advantage of the various cluster-specific gene markers recently proposed by scRNAseq studies (*6–8*) and developed the workflow presented in Fig. 1A. Thirty-three antibodies labeled with unique metal isotopes were validated (Table S1) and used to identify monocytes and macrophages (CD14, CD68, IBA1), discriminating TRMs (CD163, CD206, MERTK, LYVE1, TIM4) and IMs (HLA-DR, S100A12, SPP1, CCR2, IFI6, CLEC10A), lymphocytes (CD3, CD4, CD8, for T cells, CD20 for B cells, and CD138 for plasma cells) and stromal cells (aSMA for smooth muscle cells, CD31 for endothelial cells, both CD90 and PDPN for fibroblasts) (Fig. 1A-C and Table S1). In addition, this panel also allowed the characterization of specific cell states, such as cell proliferation (KI67) or apoptosis (cleaved-caspase3). All antibodies were validated in healthy and RA synovial tissues on the basis of their expected staining pattern, including their spatial localization in lining and perivascular areas (Fig. 1B-D and Fig. S1A). Forty high-dimensional images were analyzed for a total of 26.85mm^2^ of synovial tissue from four healthy donors, four patients with OA exhibiting a low inflammatory profile, and four patients with active RA (Disease Activity Score - Erythrocyte Sedimentation Rate (DAS28-ESR) ≥ 3.9) at baseline before the initiation of treatment. For three of the RA patients, we also included post-treatment biopsies at 6 months (Fig. S1B). Selected RA patients were positive for autoantibodies (Anti-Citrullinated Protein Antibody, ACPA^+^).

**Fig. 1.**
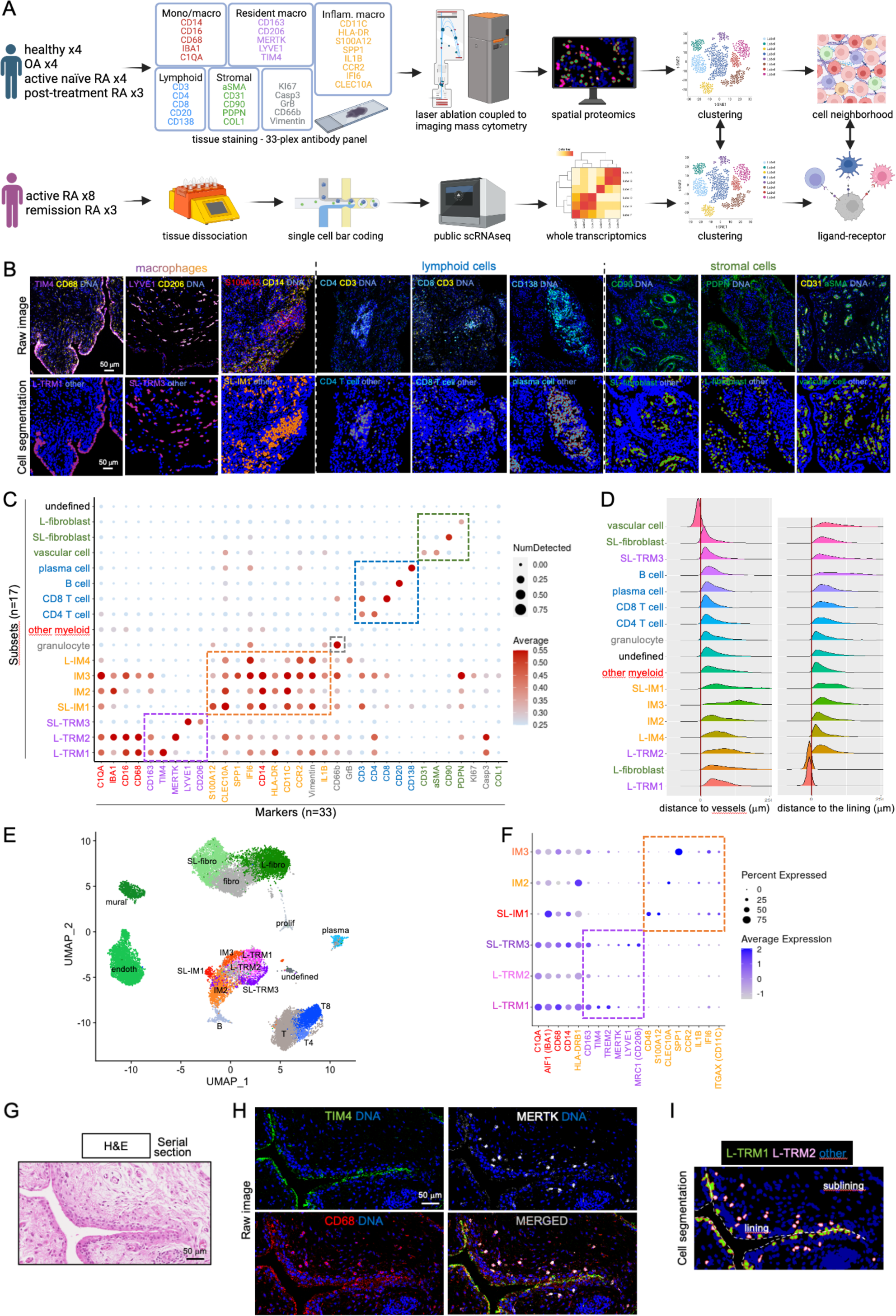
A multiplexed IMC antibodies panel to characterize the synovial stromal and immune landscape. **(A)** Schematic representation of the data acquisition workflow used for Imaging Mass Cytometry (IMC) and single cell RNA sequencing (scRNASeq). Combined analysis of spatial cell neighborhoods and ligand-receptor interactions in the different patients’ groups. Created with BioRender. **(B)** Representative pseudo-colored IMC raw images (top panels) and corresponding cluster assignments after cell segmentation (bottom panels) of the synovium samples from healthy individuals (n=4), osteoarthritis (OA, n=4) patients, early treatment-naïve rheumatoid arthritis (RA, n=4) and matched conventional synthetic (cs) disease-modifying anti-rheumatic drug (DMARD)-treated responder RA patients (n=3). Scale bar, 50μm. **(C)** Scatter plot representing the mean intensity of each marker (color code) and proportion of expressing cells (dot size) generated by Phenograph after cell segmentation of the IMC images. IM, inflammatory macrophages; TRM, tissue resident macrophages; L-, lining-; SL-; sublining-. **(D)** Distribution of each cell type in the synovium according to the minimum distance calculated to vascular cells and lining cells (L-TRM1 cells and L-fibroblasts). **(E)** UMAP plot of the different cell clusters identified from public scRNAseq datasets (*7, 15*) in the synovial tissue of RA patients with active disease (n=8, comprising 5 treatment-naïve patients and 3 DMARDs-resistant patients) or in remission (n=3, DAS28<2.6 for more than 6 months). **(F)** Scatter plot representing the mean expression of each marker (color code) and proportion of expressing cells (dot size) in the different macrophage clusters identified from scRNAseq datasets. **(G-I)** Representative H&E staining (G), IMC raw images (H) and corresponding cluster assignments after cell segmentation (I) of L-TRM1 and L-TRM2 cells in the synovium of a patient with OA. Scale bar, 50μm.

All images were then segmented into 115,411 total cells and an unsupervised clustering approach was used to classify synovial TRMs, IMs, lymphoid and stromal cells. Sixteen distinct subsets of cells were identified (Fig. 1C), and this classification was validated by comparing the raw images with the segmented/cell-subset-assigned images and confirming a clear overlap between the staining patterns (Fig. 1B and S1A). One additional undefined cell cluster was observed and did not present specific expression patterns (Fig. 1C). Importantly, by calculating the distance between each cell type from the vascular and lining cells, we obtained a comprehensive spatial cell organization allowing for the annotation of lining (L-) and sublining (SL-) cell subsets (Fig. 1D). The different cell subsets determined by IMC were also compared to the cell clusters determined using public scRNAseq datasets covering 23,759 synovial cells dissociated from the synovium of RA patients (n=11) (Fig. 1E) (*7, 15*). Using a clusterization approach based on the same markers as in previous IMC analysis, we found similar cell clusters in public scRNAseq datasets, including TRMs, IMs, T and B cells, fibroblasts, endothelial and mural cells comprising smooth muscle cells and pericytes (Fig. 1E,F and Fig. S2C). Therefore, in addition to confirming the presence of synovial single-cell clusters previously highlighted using scRNAseq (*7, 15*), IMC provides further insights into their spatial organization.

Importantly, we distinguished two distinct lining TRM cell subsets based on their differential expression of TIM4 and MERTK. Lining TRM1 cells (L-TRM1) were CD68^+^CD163^+^TIM4^+^MERTK^low^, whereas L-TRM2 cells were CD68^+^CD163^+^TIM4^-^ MERTK^high^, and both expressed the apoptosis marker cleaved-caspase3 (Fig. 1C and S2A), which is presumably linked to their capacity of phagocyting dying cells (efferocytosis) (*16*). Although L-TRM1 were located at the edge of the synovial lining layer, L-TRM2 were located slightly deeper in the tissue in the lining and proximal sublining region (Fig.1G-I). In line with previously published evidence, at the transcriptomic level, scRNAseq confirmed that L-TRM1 cells expressed higher levels of *TIM4* and *TREM2* compared to L-TRM2 cells, and both cell subsets expressed *CD163* and low levels of *MERTK* (Fig. 1F) (*7*). SL-TRM3 cells were found positive for CD206 and LYVE1, dispersed throughout the sublining membrane (Fig. 1B-F), and resembled perivascular TRM recently described in mice and RA individuals (*17*). Three distinct tissue infiltrating CD14^+^ IM clusters were defined based on varying expression of the alarmin S100A12 (SL-IM1), CD11c and HLA-DR (IM2, antigen presenting cells) or osteopontin (SPP1, IM3) (Fig.1C-F). A small proportion of IM3 cells was also positive for KI67 (Fig. 1C and S2B), suggesting cell proliferation in this cluster usually linked to tissue fibrosis (*18*). A fourth IM cluster, characterized by the expression of CCR2, IFI6 and granzyme B (L-IM4) presented an interferon signature. As expected, CD4 (helper T cells) and CD8 (cytotoxic T cells) were mutually exclusive among the CD3-expressing lymphocytes, and CD20 and CD138 respectively defined B and plasma cells. As previously described, L-fibroblasts were defined by the expression of podoplanin (PDPN^+^CD90^-^), whereas SL-fibroblasts lacked PDPN expression but exhibited CD90/THY1 expression (Fig.1B-D) (*8, 19, 20*). At the transcriptomic level, L-fibroblasts expressed *PDPN, PRG4, VCAM* and *CD55*, whereas SL-fibroblasts were characterized by the expression of *THY1, DKK3* and *CXCL12* (Fig. S2C). Due to the close proximity between those cells, both smooth muscle cells (aSMA^+^) and endothelial cells (CD31^+^) were clustered together and were identified as vascular cells (Fig. 1B-E). Overall, our results were concordant with the cell diversity described in the literature based on scRNAseq, CyTOF and multiplexed immunofluorescence technical approaches (*6, 7, 10, 17, 19–21*) but unveiled higher diversity with three TRM cell subsets displaying distinct spatial organization within the lining and sublining layers of the synovium.

### Reduced density of synovial TRMs correlates with the infiltration of immune cells in active RA synovium

Following the identification of synovial cell subsets, we compared the stromal and immune cell density across patients’ groups. A t-SNE visualization of each cell subset in the synovial tissue of healthy, OA and active treatment-naïve RA is presented in Fig. 2A, and a representative image of their spatial localization is presented in Fig. 2B-C. Their respective density (cells/mm^2^) and frequency (% of the total number of cells) in each patient group is shown in Fig. 2D-E and Fig. S3, respectively. As expected, a significant increase in both the different lymphoid cell subsets (B, plasma, CD4 and/or CD8 T cells) and the L-fibroblast cell density was observed in the synovium of RA patients as compared to healthy donors or OA patients (*19*). Although not statistically significant, we observed a trend toward an increased synovial density of IM1, IM2 and IM3 cells in active RA compared to healthy donors and their overall proportions were higher in active RA (Fig. 2D-E). Inversely, the density of most TRM, and specifically L-TRM2 cells, was significantly decreased in RA compared to OA synovial tissues. Altogether, using a precise characterization by IMC, we highlighted the reduction of all three synovial TRM cell subsets in active early treatment-naïve RA patients, concomitantly to the infiltration by numerous lymphoid and IM cells.

**Fig. 2.**
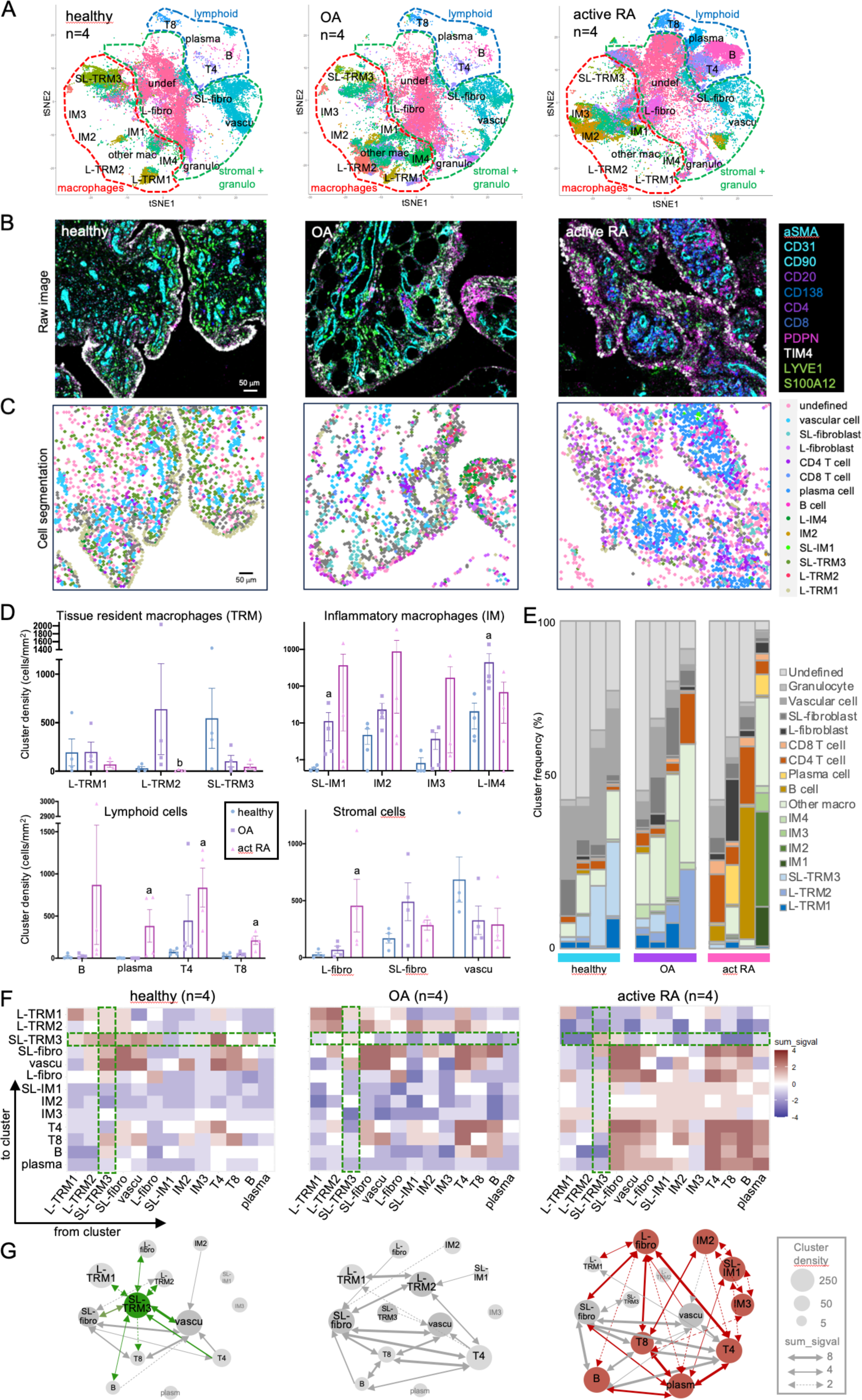
Synovial cell densities and interactions are altered in active treatment-naïve RA patients. **(A)** t-SNE plot of the different cell subsets represented in the synovial tissue of healthy individuals (n=4), osteoarthritis (OA, n=4) patients and active treatment naïve rheumatoid arthritis (active RA, n=4) patients. Macrophages (red), lymphoid (blue) and stromal (green) cell types are encircled. **(B,C)** Representative pseudo-colored IMC images of stromal (aSMA, CD31, PDPN and CD90) and immune cell markers (CD20, CD138, CD4, CD8, TIM4, LYVE1 and S100A12) of the synovium samples from healthy individuals, OA and active RA patients (B), and corresponding subset assignments after cell segmentation (C). Scale bar, 50μm. **(D)** Cell density of tissue resident macrophages (TRM), inflammatory macrophages (IM), lymphoid cells and stromal cells is presented for each patient belonging to the healthy, OA and active RA (act RA) group. Mann Whitney tests were performed: a, p<0.05 compared to healthy; b, p<0.05 compared to OA. **(E)** Cell subset frequencies in the three patients’ groups. **(F)** Heatmaps showing the cell-cell interactions as the sum of the significant values calculated for each patient (sum_sigval) belonging to the healthy (n=4), OA (n=4) and active RA (n=4) group. Brown squares indicate interactions, blue squares indicate avoidance between each cell pair. Interactions implicating SL-TRM3 cells are highlighted in green. **(G)** Schematic representation of the pairwise cell interactions, considering cluster densities (circle size) and sum_sigval values (arrow thickness) as calculated in (F). RA-specific interactions compared to both healthy and OA synovium are shown in red, interactions implicating SL-TRM3 cells that were lost in active RA synovium compared to healthy are shown in green.

### Social network analysis highlights a significant disruption of SL-TRM3 cellular interactions in active RA patients

Communications between cells in their tissue environment have been shown to dictate synovial tissue homeostasis but also RA pathogenesis through the initiation and perpetuation of inflammation, autoimmunity and tissue destruction (*7–9, 21*). To further characterize the patterns of communication between individual synovial cells, a permutation test was performed to define significant interactions between cells as compared to an empirical null distribution. Within the 5 nearest cell neighbors (k=5) and a distance threshold of 20μm, we identified interaction and avoidance behaviors between cell cluster pairs (Fig. 2F) and constructed summary graphs where nodes corresponded to cell types and each arrow matched their proximity and probability of interaction (Fig. 2G) (*22*). In healthy individuals, the main cellular network mostly involved sublining cells, including SL-TRM3, SL-fibroblasts and vascular cells. In OA patients, a similar synovial social cell network was observed, but the interactions involving SL-TRM3 cells were reduced whereas the interactions involving L-TRM2 and CD4^+^ T cells were enhanced, especially with L-TRM1 and CD8^+^ T cells (Fig. 2F, G). Importantly, in the synovial tissue of early, active, and treatment-naïve RA patients, several cell-cell interactions were disrupted compared to healthy tissues, especially the ones involving SL-TRM3 (Fig. 2F, G). Furthermore, in RA, new specific interactions were revealed, especially involving IMs, lymphoid cells and L-fibroblasts (Fig. 2F, G). These analysis therefore highlighted that the synovial infiltration by numerous inflammatory cells, including fibroblasts, lymphocytes and IMs, at early stages of RA, creates a dense cellular network in which the homeostatic social network involving SL-TRM3 is disrupted.

### The definition of spatial multicellular neighborhoods or niches reveals the dynamic and functional synovial interactions of SL-TRM3 cells

We next explored whether multicellular structural analysis, rather than pairwise cell-cell interactions, would provide further insights into the organization and dynamic changes of RA synovial tissue. We performed cellular neighborhood analysis on the IMC multiplexed imaging data and clustered the cells based on the identity of their neighbors within a radius of 20μm (*23, 24*). We identified nine cellular neighborhoods (herein called “niches”), of which three were localized in the lining (L1 to L3) and six in the sublining (SL1 to 4, ELS1 and 2) (Fig. 3A-D). Lining niches were composed of distinct proportions of L-TRM1 cells (high in L1), L-TRM2 cells (high in L2) and L-fibroblasts (high in L3). The four sublining niches (SL1-4 niches) were mostly composed of vascular cells, SL-fibroblasts and SL-TRM3 cells, which was reminiscent of the principal cell network determined in healthy donors (Fig. 2F, G). Finally, two additional niches also localized in the sublining compartment were composed of lymphoid cells and IMs cells tightly packed together, forming lymphoid aggregates, and therefore called ectopic lymphoid structures (ELS) (*5, 25, 26*). While the ELS1 niche was mostly composed of lymphoid cells, namely B cells, CD4 and CD8 T cells, the cellular composition of the ELS2 niche included B cells, plasma cells, but also the 3 IMs cell types (Fig 3A). Representative images of the niches localization are shown in Fig. 3C. Importantly, ELS1, ELS2 and L3 niches were significantly more abundant in active RA samples, compared to healthy donors or OA patients, whereas, in accordance with the disruption of the synovial lining macrophages observed in RA (*10*), the density and frequency of the L1 niche composed of L-TRM1 and 2 cells were significantly reduced early in the disease progression (Fig. 3B-C and S4). Although not statistically significant, we also observed a decreased density and frequency of the SL1 and SL3 niches, which can be partly explained by the reduced density of SL-TRM3 cells in active RA patients (Fig. 2D, E).

**Fig. 3.**
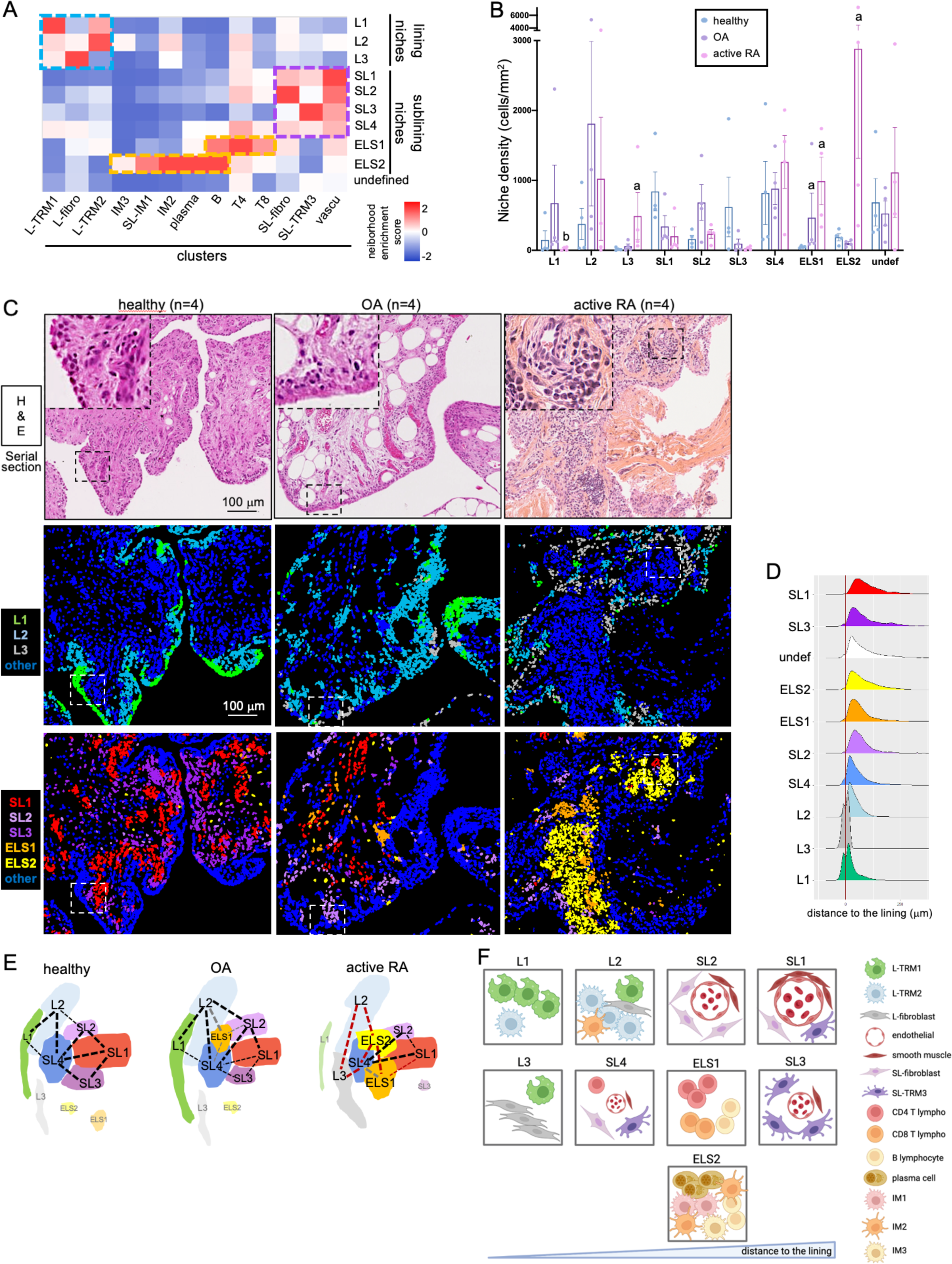
Spatial organization of the synovial niches is altered in active treatment-naïve RA patients. **(A)** Heatmap representing the neighborhood enrichment score of each cell subset across 10 niches defined in the 15 synovial tissues analyzed. Lining niches are highlighted in light blue, sublining niches in purple and ectopic lymphoid structures (ELS) niches in orange. **(B)** Niche densities are presented in the patients belonging to the healthy (n=4), osteoarthritis (OA, n=4) and active rheumatoid arthritis (active RA, n=4) group. Mann Whitney tests: a, p<0.05 compared to healthy; b, p<0.05 compared to OA. **(C)** Representative H&E staining (top panels) and corresponding images of the lining (middle panels) or sublining and ELS (bottom panels) niches in the 3 patients’ groups. **(D)** Distribution of each niche in the synovium according to the minimum distance calculated to the lining cells (L-TRM1 cells and L-fibroblasts). **(E)** Schematic representation of the niches’ spatial organization, considering the niche densities, distances to the lining and their interactions. Basal niche interactions in healthy and OA samples are provided in black and grey, additional interactions between niches observed specifically in active RA are shown in red. **(F)** Schematic representation of the cell type composition and proportions in each niche. Created with BioRender.

While each niche can represent the site of unique and local cellular interactions, distinct niches may also be interacting to support additional biological events (*24*). We therefore generated a spatial niche interaction graph for each patient group (Fig. S5A) and provided a comprehensive locoregional niches map based on the strength of the interactions between niches, their density, and their precise tissue location (Fig. 3D, E). A schematic representation of the cellular composition of each niche was also provided to summarize our findings (Fig. 3F). In healthy donors, the main interactions were observed between L1, L2, SL1, SL2, SL3, and SL4 niches. Of interest, the cells in these niches (TRMs, fibroblasts and vascular cells) are considered tissue resident or structural cells, thus highlighting the features of a basal structural and homoeostatic interaction map within the synovial tissue. In OA synovium, a similar cartography was observed but a central position of the ELS1 niche, interacting with L2, SL2 and SL4 niches, was underlined (Fig. 3E). In active RA patients, the L3, ELS1 and ELS2 synovial niches composed of all infiltrating proinflammatory immune cells and L-fibroblasts, established novel interactions between them and with the homeostatic niches, therefore highlighting profound alterations compared to the basal picture (Fig. 3E). Furthermore, we observed the loss of L1 and SL3 niches involvement in RA cellular interactive map, as well as the disruption of specific interactions, such as L2 with SL2 or with SL4 (Fig. 3E). Altogether, these results indicated that the infiltration of numerous immune cells, the formation of perivascular ELS niches, as well as the expansion of fibroblasts in the lining layer, characterizing the RA synovial tissue, have a profound impact on the whole synovial architecture and the underlying cellular communications. Most notably, as observed at the individual cell level, these results confirmed that the perivascular SL3 niche, mainly composed of SL-TRM3 cells, was profoundly altered in early active and treatment-naive RA patients.

### SL-TRM3 cells are reestablished following efficient csDMARD treatment

To study the impact of csDMARD treatment on the altered synovial cellular network observed in active RA patients, we next performed paired analysis of matched RA patients prior to therapeutic intervention and 6-months following efficient csDMARD treatment with MTX (Fig. 4A). The proportions of SL-TRM3 and vascular cells were significantly enhanced, whereas CD4 T cells and SL-fibroblasts proportions were significantly decreased by MTX (Fig. 4B). Importantly, 6-months following efficient csDMARD treatment, we observed that the cellular interactions involving SL-TRM3 cells were partially reestablished, for example with L-fibroblasts, L-TRM2 and vascular cells (Fig. 4C, D). In addition, MTX treatment induced novel interactions, such as between SL-TRM3 and plasma cells, whereas preexisting interactions persisted post-treatment, such as with SL-fibroblasts and CD4 T cells (Fig. 4C, D). Interestingly, upon MTX treatment and along with an increased proportion of the SL1 and SL3 niches (Fig. 4E and S4), SL3 reestablished significant connections with SL1 and SL4 (Fig. 4F, G and S5B), as observed in homeostatic conditions (Fig. 3E). Similarly, the interaction between L2 and SL4 niches was recovered in the synovial tissue from responder RA patients, and the strength of several interactions, such as ELS2-L2, L2-L3 or SL4-L3, was downregulated compared to baseline RA (Fig. 4F, G). Overall, the response to csDMARD therapy was associated with a partial recovery of the basal homeostatic synovial cartography, sustaining the particular involvement of SL-TRM3 cells. In particular, we highlighted the key role of SL-TRM3 intra- and inter-niche interactions with vascular cells and SL-fibroblasts (SL1, SL3 and SL4 niches), CD4+ T and plasma cells (ELS1 and ELS2 niches), L-fibroblasts and L-TRM2 cells (L2 niches) in treatment response.

**Fig. 4.**
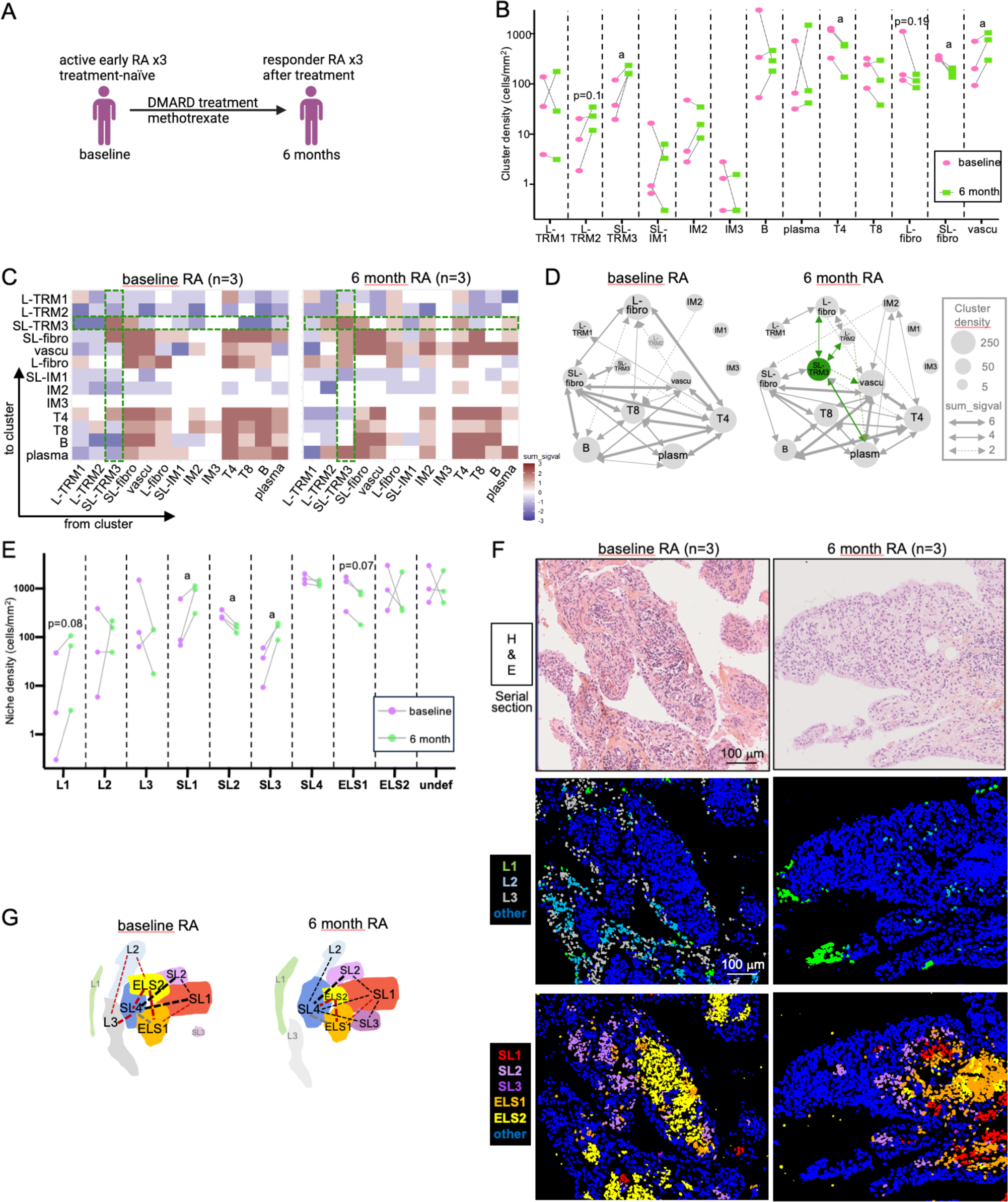
Partial recovery of the synovial architecture following efficient csDMARD treatment. **(A)** Synovium samples were obtained from active treatment-naïve RA patients at baseline (n=3) and 6 months following efficient treatment with the csDMARD methotrexate (MTX) from the same patients (n=3). **(B)** Paired analysis of cell subset densities in RA patients at baseline and 6 months post-csDMARD treatment. Paired student’s t-tests were performed: a, p<0.05. **(C)** Heatmaps showing the cell-cell interactions as the sum of the significant values calculated for each patient (sum_sigval) belonging to the active RA at baseline (n=3) and 6 months post-csDMARD treatment (n=3) group. Brown squares indicate interactions, blue squares indicate avoidance between each cell pair. Interactions implicating SL-TRM3 cells are highlighted in green. **(D)** Schematic representation of the pairwise cell interactions, considering cluster densities (circle size) and sum_sigval values (arrow thickness) as calculated in (C). Interactions implicating SL-TRM3 cells that were induced 6 months post-csDMARD treatment compared to baseline are shown in green. **(E)** Paired analysis of niche densities in RA patients at baseline and 6 months post-csDMARD treatment. Paired student’s t-tests were performed: a, p<0.05. **(F)** Representative H&E staining (top panels) and corresponding images of the lining (middle panels) or sublining and ELS (bottom panels) niches in the synovium of RA patients at baseline and 6 months post-csDMARD treatment. **(G)** Schematic representation of the niches’ spatial organization, considering the niche densities, distances to the lining and their interactions. Basal niche interactions previously observed in healthy and OA samples are provided in black and grey, additional interactions between niches observed specifically in RA samples are shown in red (refers to Fig. 3E).

### Combined spatial synovial mapping and scRNAseq ligand-receptor analysis underline SL-TRM3 cellular pathways linked to csDMARD treatment response

To further highlight the pathways that govern cell communications within the synovial niches, we further analyzed the public scRNAseq datasets of synovial cells dissociated from the synovium of patients with active RA (n=8) and in remission (n=3) introduced previously (Fig. 1E,F) (*7, 15*). We compared the differential ligand-receptor interactions between active and remission RA samples (Fig. S6). Interestingly, several ligand-receptor interactions were induced in the synovial tissue of RA patients in remission when compared to active RA, such as interactions involving SL-TRM3 (Fig. 5A), in line with the spatial analysis presented above (Fig. 4). Indeed, SL-TRM3 cells from remission RA patients were found, for instance, to be major producers of *IL10, IL1B* and complement *C3*, which target IMs and TRMs (Fig. 5B, C). In addition, the expression of *CLEC2B* and *CD86* by SL-TRM3 was increased and these molecules signal to CD4 and CD8 T cells through *KLRB1* and *CD28* respectively (Fig. 5B, D). SL-TRM3 cells from RA patients in remission produced significantly more *TWEAK (TNFSF12)* and *IGF1* directed to the vascular cells and SL-fibroblasts (Fig. 5B, E) compared to active RA. Importantly, we also highlighted that SL-TRM3 cells expressed a high level of *GAS6* to stimulate L-TRM1 and 2 through its receptor *MERTK*, but also IM2, mural cells, SL- and L-fibroblasts through its other receptor *AXL* (Fig. 5B, C, E).

**Fig. 5.**
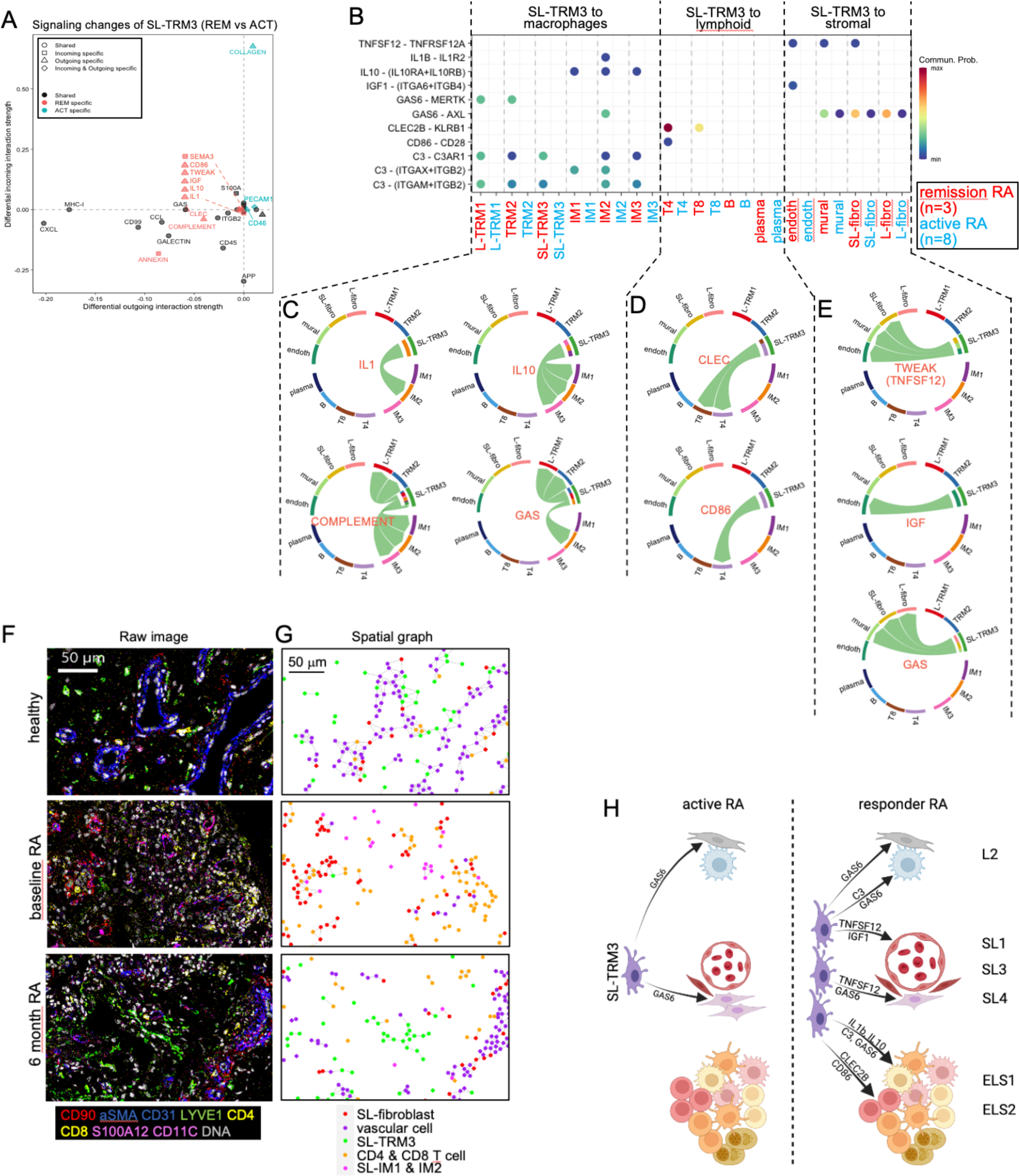
Combined IMC and scRNAseq ligand-receptor analysis underlines the existence of cellular pathways linked to DMARDs treatment response in SL-TRM3 cells. Public scRNAseq datasets (*7, 15*) were used to analyze the potent ligand-receptor interactions between SL-TRM3 cells and other synovial cells in the synovium of RA patients with active disease (n=8, comprising 5 treatment-naïve patients and 3 DMARDs-resistant patients) or in remission (n=3, DAS28<2.6 for more than 6 months). **(A)** Dot plot of all the differential outgoing and incoming interactions involving SL-TRM3 cells in active RA (ACT) versus remission RA (REM) samples. Active RA-specific interactions are shown in blue, remission RA-specific interactions in red. **(B)** Examples of remission RA-specific interactions involving SL-TRM3 cells as sender cells are provided as bubble plots. **(C-E)** Examples of remission RA-specific interactions are provided as chord diagrams. SL-TRM3 sender cells are presented at the start of the arrows and receiver cells at the end. Interactions from SL-TRM3 cells to macrophages are provided in (C), to lymphoid cells in (D) and to stromal cells in (E). **(F)** Representative Imaging Mass Cytometry (IMC) raw images in the synovium samples from healthy individuals (n=4) and RA patients at baseline (n=3) and 6 months post-csDMARD treatment (n=3) showing SL-fibroblasts (CD90^+^), vascular cells (aSMA^+^ or CD31^+^), SL-TRM3 cells (LYVE1^+^), T cells (CD4^+^ or CD8^+^), SL-IM1 cells (S100A12^+^) and IM2 cells (CD11C^+^). **(G)** Corresponding spatial graphs presenting the interactions (grey nodes) between SL-fibroblasts (red), vascular (purple), SL-TRM3 (green), T (orange), SL-IM1 and IM2 (pink) cells in the 3 patients’ groups. **(H)** Schematic representation of the differential cell-cell communications and implicated molecular pathways established by SL-TRM3 cells in the synovium of active versus responder RA patients. Location of the cells in their respective niches is also provided. Created with BioRender.

The results obtained from these differential ligand-receptor interactions were supported by our spatial analysis highlighting, among others, significant connections (i) between SL-TRM3, vascular cells and SL-fibroblasts (within SL1, SL3 and SL4 niches), (ii) between SL-TRM3, L-fibroblasts and L-TRM2 cells (within SL4 and L2 niches) or (iii) between SL-TRM3 and CD4 T cells (within SL4 and ELS1 niches) in responder RA patients (Fig. 4). In situ, these cellular interactions were visualized using our IMC analysis and both raw images of LYVE^+^ SL-TRM3 cells in their tissue microenvironment and the corresponding spatial graphs highlighted clear proximities between SL-TRM3, SL-fibroblasts, vascular cells, T cells, SL-IM1 and IM2 cells, especially in RA patients 6-months following csDMARD treatment (Fig. 5F, G). A schematic representation of these differential cellular interactions and pathways in active versus responder RA patients is provided in Fig. 5H.

## DISCUSSION

Here, we provided an unprecedent comprehensive synovial cell and niche social network mapping by performing multiplex, single-cell spatial analysis of the synovial tissue using IMC. This approach allowed to identify and decipher the spatial organization of the main tissue resident / homeostatic or infiltrating / inflammatory myeloid, lymphoid, and stromal cells. We compared, for the first time, healthy tissues with OA and matched active treatment-naïve and csDMARD-treated responder RA synovium and highlighted disease- and treatment-response-specific alterations of the synovial cellular network and their underlying disrupted cellular pathways compared to homeostatic conditions.

Using a 33-antibodies panel, we were able to distinguish sixteen distinct immune and stromal cell subsets and to spatially map each of these subsets within the lining and the sublining synovial regions. Overall, we highlighted the presence of seven macrophage subsets, for which the spatial features and expression profile confirmed and expanded the diversity described in the literature using scRNAseq (*6, 7, 11, 21*). By using a distinct but complementary approach and clustering the cells based on the identity of their neighbors within the synovium, we then identified the presence of nine relevant synovial niches, of which three were localized in the lining (L1 to L3) and six in the sublining (SL1 to 4, ELS1 and 2).

We first underlined that two distinct TRM cell subsets can be found in the lining layer, depending on their expression of TIM4 and MERTK. L-TRM1 cells expressed *CD163*, *TIM4, TREM2* and *MERTK*, were located at the edge of the lining layer and were abundant in healthy synovium, in line with their previously described homeostatic, phagocytic and anti-inflammatory functions (*7, 11*). L-TRM2 cells expressed lower level of *TIM4* and *TREM2*, correlated to MERTK^+^TREM2^low^ cells described previously (*7*). We show here that they can be found deeper in the proximal sublining layer and were most abundant in OA synovium. Whether L-TRM2 cells represent the pro-inflammatory state of L-TRM1 cells with lost barrier and homeostatic functions or originates from another cell subset now deserves further studies (*7, 11*). We also observed that lining fibroblasts expressing PDPN were expanded in RA and mainly localized in the L3 niches, where, as previously described in the literature (*8, 19, 21*), they assume a central position by connecting with lining L-TRM1 cells but also with sublining cells and niches. This expansion matched with a significant reduction of the density and frequency of the L1 niche mainly composed of L-TRM1 cells in RA, confirming previous studies that demonstrated a significant disruption of the lining myeloid barrier that physically secludes the joint in homeostatic conditions (*4, 7, 10, 11*).

Within the sublining compartment, notably, different subsets of IMs and lymphoid cells were shown to communicate together and form two distinct ELS niches. Such ELS organization recapitulates the architecture of secondary lymphoid organs, with (i) a distinct T cell rich zone in contact with a B cell rich area; (ii) a network of follicular dendritic cells (FDC) and activated resident stromal cells such as fibroblasts; and (iii) the presence of high endothelial venules (HEVs) that participate to T and B cells recruitment and in situ differentiation into plasma cells, that produce somatically mutated disease-specific autoreactive antibodies (*27–29*). Our multiplexed spatial approach supports this view and provides additional insights into ELS organization. The ELS1 niche has a relative enrichment for T cells and few B cells, corresponding with the T cell rich zone, whereas the ELS2 niche has many more B cells and fewer T cells, corresponding with the B cell rich area. Of note, CD11C^+^HLADR^+^ myeloid cells (IM2 cells), resembling FDC, were observed withing the ELS2 niche. In the context of active RA and before any treatment intervention, both ELS niches were also strongly connected with resident sublining fibroblasts (PDPN^-^CD90^+^) near vascular vessels belonging to the SL1 and SL4 niches, consistent with the described presence of activated stromal cells and HEVs, respectively (*27–29*). Importantly, we also highlighted the presence of different subtypes of IMs cells within the ELS2 niche. Although analyzing the expression of additional markers would be necessary to further characterize ELS cells such as FDC, T follicular helper (Tfh) cells, T regulatory (Treg) cells, plasmablasts, activated stromal cells, and endothelial cells, our data clearly indicates that several inflammatory myeloid cells with potent roles on monocyte and fibroblast activation (IM1, S100A12^+^), antigen presentation (IM2, HLA-DR^+^), neutrophil activation and matrix remodeling (IM3, SPP1^+^) (*6, 7, 11, 30–32*) were localized in closed proximity to the cells involved in ELS functions and enriched in active RA.

As despite the availability of a large variety of conventional, biologic, and synthetic DMARDs approved for the treatment of RA, 40% of patients still do not respond to treatment, understanding molecular signatures underlying the variety of clinical and treatment-response phenotypes remains one of the major areas of research in rheumatology, with the aim of developing personalized therapies for RA patients. Here, we chose to focus our analyses on RA patients who, 6-months following csDMARD treatment, presented a good or moderate treatment response, in order to decipher the complex biological mechanisms driving the resolution of inflammation. We demonstrated that a key cellular social network involved in the homeostasis of healthy synovium comprised SL-TRM3 cells, SL-fibroblasts and vascular cells in SL1 to 4 niches within the sublining membrane, while, in active RA, the loss of SL-TRM3 cells induced profound changes in the SL1 and SL3 organization. Importantly, csDMARD treatment was able to induce a partial recovery of the basal homeostatic synovial cartography i.e. part restoration of SL-TRM3 cells, allowing, for example, the reorganization of lining fibroblasts and TRMs cells similar to the healthy or OA situation. The homeostatic proportion of SL-TRM3 was indeed partly recovered 6-months after MTX treatment, and these cells established strong interactions with IMs, TRMs, lymphoid and stromal cells. Perivascular SL-TRM3 cells are positive for the hyaluronan receptor LYVE1 and the mannose receptor CD206, and are thought to control homeostatic collagen production, prevent leukocyte infiltration and fibrosis (*11, 33, 34*). They are therefore well adapted to exert beneficial roles in response to treatment.

To overcome the technical limitation that currently restricts the maximum number of markers that can be simultaneously analyzed using the IMC approach (33 antibodies in this study), we further dissected the molecular basis of synovial cell communications by combining IMC with public scRNAseq datasets analysis and explored a large number of cell-cell communications based on ligand-receptor interactions. We observed that in the synovial tissue of RA patients in remission, SL-TRM3 cells communicate with IMs and TRMs through *IL10* and complement *C3*, with the presumed scope of limiting inflammation, cytokines production and stimulate the phagocytosis of damaged materials and immune complexes (*35–37*). SL-TRM3 cells also produced more *TWEAK (TNFSF12)* and *IGF1* targeting vascular cells and SL-fibroblasts, supporting the notion that these pathways are linked to remission (*38–40*). In addition, DMARD treatment significantly influenced SL-TRM3 to express more *CLEC2B* and *CD86* to interact with *KLRB1*^+^ and *CD28*^+^ T cells, thus also regulating T cell functions (*41, 42*). Importantly, upon efficient treatment, SL-TRM3 cells expressed high levels of *GAS6* ligand to stimulate stromal cells and macrophages upon binding to its receptors, *AXL* and *MERTK*. TAM (TYRO3, AXL, and MERTK) receptors have been shown to negatively regulate the inflammatory cascade involved in RA pathophysiology and mediate phagocytosis of apoptotic cells (*43*). These findings, supported by our spatial cell social networks and synovial niche mapping, sustain the development of TAM-targeted therapeutic strategies to foster the resolution of inflammation in the context of arthritis (*7, 11, 44–46*). Overall, our combined spatial and scRNAseq analysis contribute new clues to decipher the complexity involved in synovial homeostasis and response to DMARD therapy.

Our findings must be considered in the context of some limitations, including the relatively low number of samples analyzed by IMC. The synovial tissues from four ACPA positive RA patients were analyzed, selected to have the most homogeneous synovial histopathological profile, all being characterized by a lympho-myeloid pathotype. Further studies should assess whether tissues presenting other histopathological profiles (pauci-immune and diffuse-myeloid (*5*) exhibit specific disease- and treatment-response related cellular and molecular pathways. Moreover, although one of the strengths of our study consist in the comparison of the expression of 33 proteins between paired synovial samples of active treatment-naïve and csDMARD-treated responders RA patients, our combined scRNAseq analysis based on public datasets focused on the comparison between active and remission RA patients. In accordance with the IMC data, the scRNAseq analysis validated our key findings in an independent cohort, further supporting the robustness of out findings.

In summary, we provided an unprecedented single-cell profiling of the synovial tissue in homeostatic conditions and in the context of RA before and after csDMARD treatment. Combining transcriptomic and proteomic approaches, we highlighted several pathways of DMARDs response based on the elucidation of the interconnections between the stromal microenvironment and the immune cell landscape. Overall, this study forms the basis for the development of more targeted, hence more active therapeutic strategies for RA.

## MATERIALS AND METHODS

### Synovial tissue samples for IMC

Synovial tissues were obtained by ultrasound-guided synovial biopsy from DMARD-naïve patients with early and active RA (n = 4), and from the same patients 6-months following the initiation of MTX treatment (n = 3) (Fig. S1B). Patients were recruited in the Pathobiology of Early Arthritis Cohort (PEAC) cohort (http://www.peac-mrc.mds.qmul.ac.uk) and enrolled at Bart’s Health NHS Trust in London as previously described (*47*). All patients fulfilled the 2010 EULAR criteria for RA (*3*). Patients had active disease with duration of symptoms <12 months and were all naïve of treatment (corticosteroids, csDMARDs or biologic therapies). All procedures were performed following written informed consent and were approved by the hospital’s ethics committee (REC 05/Q0703/198). Patients selected for this study were positive for autoantibodies (Anti-Citrullinated Protein Antibody, ACPA^+^), and their synovial tissue presented an infiltration of macrophages and lymphoid cells (T, B and plasma cells) and were classified as belonging to the lympho-myeloid pathotype (*5*). After 6-months of treatment, patients were either good (delta DAS28-ESR ≥ 3.4, n = 3) or moderate (delta DAS28-ESR = 2.8, n = 1) responders (Fig. S1B).

Synovial tissues from OA patients (n=4) were obtained at the time of total joint replacement. Enrolled OA patients were diagnosed according to EULAR recommendations (*48*), were identified as having no overlapping diagnosis with any other type of rheumatic disease, underwent knee arthroplasty and were included without other prior selection (Fig. S1B). Healthy synovial tissues from patients undergoing meniscal or cruciate ligaments surgery presenting a non-inflamed synovium (assessed via magnetic resonance imaging, macroscopically and histologically) (n=4) were used as control. OA and control biopsies were obtained following written patient’s informed consent and ethics approval was granted by the Nantes University Hospital ethics committee (DC-2011-1399).

All biopsies were fixed in 4% formaldehyde, embedded in paraffin and underwent Haematoxyline and Eosin (H&E) staining to confirm the quality of the tissue and the presence of typical synovial features, a lining and a sublining, as well as ectopic lymphoid structures for RA selected tissues (Fig. S1B).

### Antibody validation and metal conjugation

A panel of 33 antibodies targeting the proteins listed in Table S1 was validated prior to the IMC analysis. Except for already validated and coupled antibodies provided by Fluidigm or previously used by our IMC facility, all other antibodies were titrated and validated on synovial tissues from RA patients and human tonsil as control by immunofluorescence (IF) experiments (see supplementary materials section). Once the ideal titer was determined, antibodies were coupled to heavy metal using the X8 Antibody Labeling Kit (Standard Biotools) according to the manufacturer’s instructions at the CIM facility. To definitively validate the panel of 33 antibodies and their different titers, we performed preliminary IMC staining on RA synovium and confirmed the expected staining features according to our experience and the literature.

### Tissue Labeling for Imaging Mass Cytometry

Two serial 3µm tissue sections were placed on Polysine-treated microscope slides. The first slide was stained with H&E for selecting the ROIs. The second one was used for metal-tagged antibodies labeling and IMC ablation. For each section, a maximum of 2 mm² of tissue including a lining and a sublining layer were chosen according to the H&E staining.

After deparaffinization and antigen retrieval using Dako Target Retrieval Solution at pH 9 (S236784-2, Agilent technologies) in a water bath (96°C for 30 min), tissue sections were delimited with a hydrophobic pen, incubated with Superblock™ (37515, ThermoFisher Scientific) at room temperature (RT) for 45 min, and then with FcR Blocking Reagent (130-092-575, Miltenyi) at RT for 1h. After washes in PBS/0.2% Triton X-100 (PBS-T), the cocktail of metal tagged antibodies (Table S1) was prepared in PBS/1% BSA buffer. Following the incubation with the primary antibodies at 4°C overnight, the sections were washed in PBS-T and nuclei were stained with iridium (1:400 in PBS; Fluidigm, Standard Biotools), a DNA Intercalator, for 30 min at RT. Sections were then washed in PBS, then in distilled water, and finally dried at RT for 30 min.

### Imaging Mass Cytometry acquisition

Images were acquired with the Hyperion Imaging System (Fluidigm, Standard Biotools) according to the manufacturer’s instructions. After choosing the ROI in the section, the ROI was ablated with a UV laser at 200Hz. Data were exported as MCD files and visualized using the Fluidigm MCD™ viewer v1.0.560.6. The minimum and maximum thresholds were adapted for each marker and for each tissue for optimal visualization.

### Data analysis

Before running the unsupervised analysis, the quality of the staining for each marker on each image was qualitatively controlled, and forty high-dimensional images for a total of 26.85mm^2^ of tissue were analyzed.

#### Cell segmentation

Cell segmentation was performed as previously described (*49*). Briefly, pixel classification and training were performed using Ilastik v1.3.3 to generate a probability map. A segmentation mask was then created from the probability map using CellProfiler v4.1.3.

#### Cell Clustering

The cells were clustered using the Phenograph algorithm (v0.99.1) in Rstudio (v4.2.0) after a normalization of the 99^th^ percentile, the outliers were eliminated, and the background reduced. The following markers were used for a first clustering, with a k-value set to 30: CD16, CD20, CD31, CD66b, CD68, CD90, CD138, CD163, CD206, aSMA, CLEC10A, HLADR, IBA1, IFI6, C1QA, IL1B, LYVE1, MERTK, SPP1, PDPN, S100A12, TIM4, Vimentin, CCR2, CD3, CD4, CD8, CD11C, CD14. To better discriminate the different sub-populations, we then sub-clustered monocytes / macrophages expressing CD14, CD68, IBA1, CD163 and/or CD206 using CD16, CD68, CD163, CD206, CLEC10A, HLADR, IBA1, IFI6, C1QA, IL1B, LYVE1, MERTK, SPP1, S100A12, TIM4, CCR2, CD11C, CD14, with a k-value set to 140. The non-macrophage clusters were sub-clustered using the expression of CD20, CD31, CD66b, CD90, CD138, aSMA, PDPN, Vimentin, CD3, CD4, CD8 with a k-value set to 140. Finally, similar sub-clusters were merged, annotated and the macrophages and non-macrophages populations were reunited in a single data frame for further analyses.

A t-SNE (Barnes-Hut t-SNE method) dimensional reduction was performed to visualize the data on 2D maps, after a concatenation and a 99^th^ percentile normalization of the data (initial dimensions = 110; perplexity = 30; theta = 0.5; max iterations = 1000). Scatter plots of the mean intensity for each marker and proportion of expressing cells were generated using the ggplot2 package (v3.4.0).

#### Distances and cell-cell interactions

Each cell was assigned a minimum distance value from either vascular cells or lining cells (L-TRM1 cells and L-fibroblasts) using the minDistToCells function of the R package imcRtools (v1.5.2) (*50*). Data were represented using the ggplot2 package (v3.4.0).

Cell-cell interactions were determined and visualized using the R packages imcRtools (v1.5.2) and cytomapper (v1.8.0) (*51*), respectively. A permutation test was performed to define the positive or negative interactions with neighboring cells. Cells were considered as neighbors within the 5 nearest cell neighbors (k=5) and a distance threshold of 20 μm, using the Histocat knn interaction graph method. The plotSpatial function was used to visualize cells’ centroids and cell-cell interactions. For each patient, a statistical analysis of these pairwise cell interactions was performed using the testInteractions function which provided significant values of 1 in case of cell-cell interaction (sigval =1, p <0.01) or −1 in case of avoidance (sigval = −1, p <0.01). The results were visualized by computing the sum of the sigval (sum_sigval) using ggplot2.

#### Cellular neighborhood analysis

Cellular neighborhood (niche) analysis was performed as described earlier by Schurch et al. (*22*), with the expansion graph method set to 20 μm. A final number of 10 niches (kmeans centers = 10) was chosen based on their location in the tissue and their cell type enrichment score. To describe tissue regions in which niches may be interacting, we analyzed their spatial organization as described in (*23*) using the detectSpatialContext function with the expansion graph method and a threshold of 0.9. We then filtered all the niche interactions with a minimum of 3 patients and 500 cells involved. The plotSpatialContext function was used to visualize niche interactions in the different patients’ groups. Schematic representations of the niches spatial features were created with BioRender.

### Single-cell RNA-seq analysis

Preprocessed scRNA-seq count data were retrieved from two public datasets (E-MTAB-8322 (*7*), GSE200815 (*15*)) covering single synovial cell data from patients with active RA (n=8) and remission RA (n=3). Active RA patients included both treatment-naïve patients (n=5) and DMARDs-resistant patients (n=3), the latter being treated for more than 6 months and resistant to both MTX and TNF inhibitors. Cells characterized by the expression of less than 200 genes or more than 20% mitochondrial genes were removed. Count data were SCTransformed using Seurat (v4.2.0). We then corrected sample-driven batch effects with the Harmony (v0.1.0) function and projected the cells into two dimensions with UMAP using 50 principal components. Seurat was used to identify clusters with a resolution setting of 0.6. Macrophages and T cells were sub-clustered with a resolution setting of 0.3. The clusters were manually annotated according to IMC clusters by identifying the differentially expressed genes using the FindAllMarkers function in Seurat and focusing on the markers used for IMC. Clusters annotated as “fibro”, “prolif”, “undefined” and “T cells” in Fig. 1E were not considered for further analyses. Scatter plots of the mean expression for each marker and proportion of expressing cells were generated using ggplot2.

Ligand-receptor analysis were performed using the CellChat (v1.6.1) R package workflow using default settings, adding the S100As ligand-receptor interactions (*52*) into the CellChatDB interaction file. We compared active RA versus remission RA samples. All outgoing / incoming signaling patterns and differential interactions were first computed. We then focused on all signaling changes of SL-TRM3 cells in active versus remission RA samples. For each analysis, we provided examples of differential ligand-receptor interaction using the netVisual chord diagram and bubble plot functions.

### Statistics

Statistical analyses were performed using GraphPad Prism v8 statistical software. P-values of <0.05 were considered significant and data were expressed as mean ± standard error of mean (SEM). Differences in quantitative variables were evaluated by the Mann–Whitney U test. Paired Student’s t-test was used to perform paired-analysis of cell frequencies at baseline and 6 months after treatment.

The investigators were blinded to sample group allocation during experiments and analysis.

## Acknowledgements

We are thankful to the MicroPICell facility for advice on IF and confocal microscopy (S. Nedellec and P. Paul-Gilloteaux), the “Institut de la Main Nantes Atlantique” (L. Ardouin), and the department of orthopedic surgery of the Nantes University Hospital (D. Waast) for providing biopsies. The authors also acknowledge the SC3M plateform from the Inserm/NU/ONIRIS UMR1229 RMeS Laboratory and SFR François Bonamy-UMS 016 for their help to manage bio-informatic datasets, biopsies, and prepare slides (J. Veziers, Y. Le Guennec and M. Dutilleul).

## Author contributions

The following author contributions are noted: GC, HAM, CP, FA, FB study conceptualization and funding acquisition; JDL, YG, ASD, ALP histology and IMC acquisition; JDL, LAC, OB, FB IMC bioinformatic analysis; OB, FB scRNAseq analysis; LF, CP, HAM, BLG provided additional materials and resources relative to patient samples and IMC antibodies; JDL, LAC, OB, FB data visualization and figure generation; GC, AN, MAB, JG, CP, FA, FB writing of the original draft, review and editing.

## Data and materials availability

The raw data and analysis codes supporting the conclusions of this article will be made available by the authors, upon reasonable request.

## Supplementary Materials

### Antibody validation by immunofluorescence

3µm-thick tissue sections of interest were placed on a Polysine-treated microscopy slide (P4981001, Thermo Fisher Scientific). The sections were deparaffined and a step of Heat-Induced Epitope Retrieval in a Tris-EDTA solution pH=9 for 20 hours at 60°C was performed. The slides were then rinsed in Tris-Buffered Saline-Tween 20 (TBS-T, F/TBT010, Microm Microtech France) solution and non-specific binding sites were blocked with 5% Normal Donkey Serum (017-000-121, Jackson ImmunoResearch) during 30 minutes at room temperature in a moisture chamber. Primary antibodies (Table S1) were then incubated overnight at 4°C. Following rinsing, secondary antibodies were diluted at 1:500 in blocking solution and incubated during 1 hour at room temperature. All the secondary antibodies were coming from Jackson ImmunoResearch, raised in Donkey and coupled to a fluorochrome. The nuclei were then stained with DAPI (D9542, Sigma-Aldrich) diluted at 1:2000 during 10 minutes at room temperature. Finally, slides were mounted in ProLong® Gold Antifade Reagent (P36930, Invitrogen) and imaged using a Nikon A1rHD LFOV confocal microscope.

