## Supplementary data for "Spatial mapping of rheumatoid arthritis synovial niches reveals specific macrophage networks associated with response to therapy"

### Table S1. Antibody panel for IMC

| Target | Clone | Supplier | Metal | Dilution |
|---|---|---|---|---|
| **Collagen type 1** | Polyclonal | Fluidigm | 169 Tm | 1/500 |
| **Alpha-SMA** | 1A4 | Fluidigm | 141 Pr | 1/2000 |
| **CD31** | EPR3094 | Fluidigm | 151 Eu | 1/200 |
| **Vimentin** | D21H3 | Fluidigm | 143 Nd | 1/2000 |
| **CD90** | Polyclonal | R&D Systems | 148 Nd | 1/100 |
| **PDPN** | NZ-1.3 | Thermo Fisher | 164 Dy | 1/100 |
| **CD66b** | REA306 | Miltenyi | 89 Y | 1/100 |
| **CD16** | EPR16784 | Fluidigm | 146 Nd | 1/800 |
| **CD14** | EPR3653 | Fluidigm | 144 Nd | 1/400 |
| **CD68** | KP1 | Fluidigm | 159 Tb | 1/2000 |
| **IBA1** | EPR16588 | Abcam | 153 Eu | 1/200 |
| **CD163** | EDHu-1 | Fluidigm | 147 Sm | 1/400 |
| **CD206** | 685645 | R&D Systems | 176 Yb | 1/100 |
| **MERTK** | Y323 | Abcam | 173 Yb | 1/500 |
| **LYVE1** | Polyclonal | Abcam | 145 Nd | 1/50 |
| **TIM4** | Polyclonal | R&D Systems | 171Er | 1/200 |
| **HLA-DR** | TAL-1B5 | Abcam | 174 Yb | 1/2000 |
| **S100A12** | Polyclonal | Abcam | 158 Gd | 1/100 |
| **SPP1** | 7C5H12 | Abcam | 170 Er | 1/500 |
| **CCR2** | 48607 | R&D Systems | 165 Ho | 1/50 |
| **IFI6** | Polyclonal | Abcam | 166 Er | 1/50 |
| **CLEC10A** | Polyclonal | Abcam | 149 Sm | 1/200 |
| **IL1B** | 3A6 | CST | 160 Gd | 1/200 |
| **C1QA** | EPR2980Y | Abcam | 161 Dy | 1/200 |
| **CD11C** | EP1347Y | Abcam | 154 Sm | 1/200 |
| **CD3** | D7A6E | CST | 152Sm | 1/200 |
| **CD4** | EPR6855 | Fluidigm | 156 Gd | 1/100 |
| **CD8a** | C8/144B | Fluidigm | 162 Dy | 1/200 |
| **CD20** | EP459Y | Abcam | 142 Nd | 1/200 |
| **CD138** | MI15 | Stem Cell | 150 Nd | 1/200 |
| **Ki-67** | B56 | Fluidigm | 168 Er | 1/800 |
| **Granzyme-B** | EPR20129-217 | Fluidigm | 167 Er | 1/1000 |
| **cleaved caspase-3** | 5A1E | Fluidigm | 172 Yb | 1/400 |
| **Iridium** | - | Fluidigm | 191/193 Ir | 1/400 |

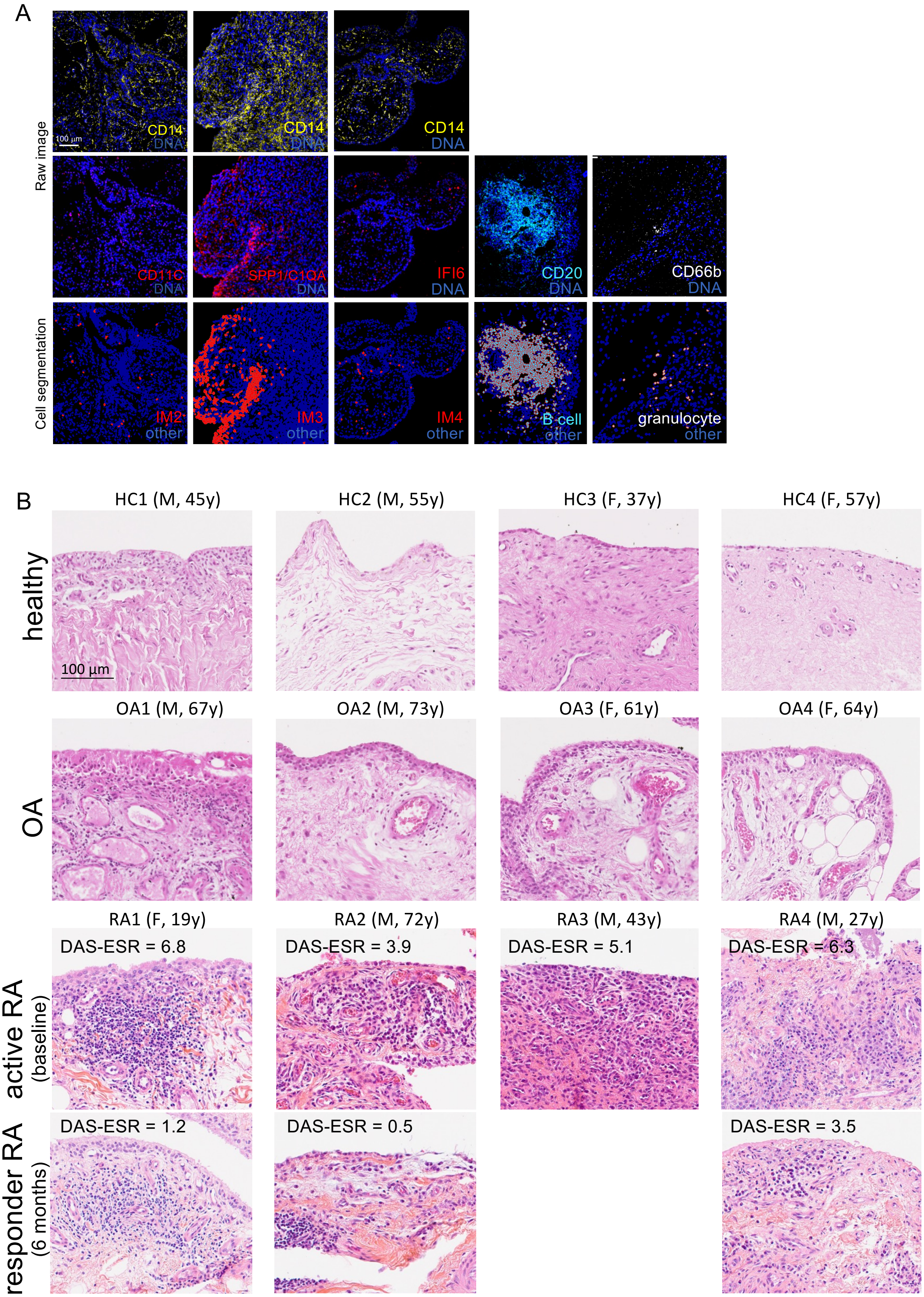


**Fig. S1. Patients' sample and validation of IMC cell clusters.**
**(A)** IMC raw images (top panels) and corresponding cluster assignments after cell segmentation (bottom panels). Refers to Fig. 1B. Scale bar, 100µm. **(B)** H&E staining of all samples from healthy controls, OA and RA patients. Gender and age is provided, as well as the DAS-ESR for RA patients. Patients RA1-3 are considered good responders to methotrexate treatment after 6 months (delta DAS-ESR ≥ 3.4). The RA4 patient is considered a moderate responder (delta DAS28-ESR = 2.8). Sample of RA3 patient after 6 months was not analyzed.

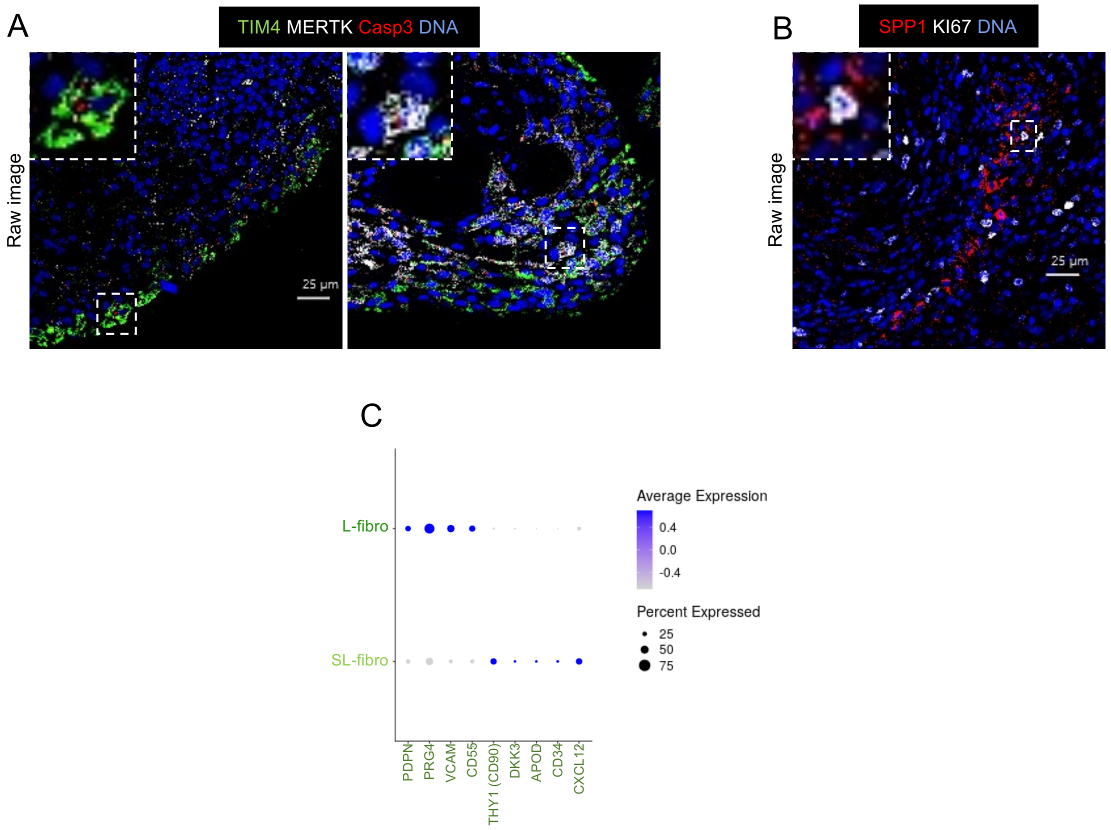


**Fig. S2. Characterization of cell subsets by IMC and scRNAseq analysis.**
**(A)** IMC raw images showing positive staining for active caspase-3 (red) in TIM4$^{+}$MERTK$^{low}$ (L-TRM1 cells, left panel) and TIM4$^{low}$MERTK$^{high}$ (L-TRM2 cells, right panel) cells. Scale bar, 25µm. **(B)** IMC raw images showing positive staining for KI67 (white) in SPP1$^{+}$ cells (IM3 cells). Scale bar, 25µm. **(C)** Scatter plot representing the mean expression of each marker (color code) and proportion of expressing cells (dot size) in the two fibroblast clusters identified from public scRNAseq datasets (refers to Fig. 1E).

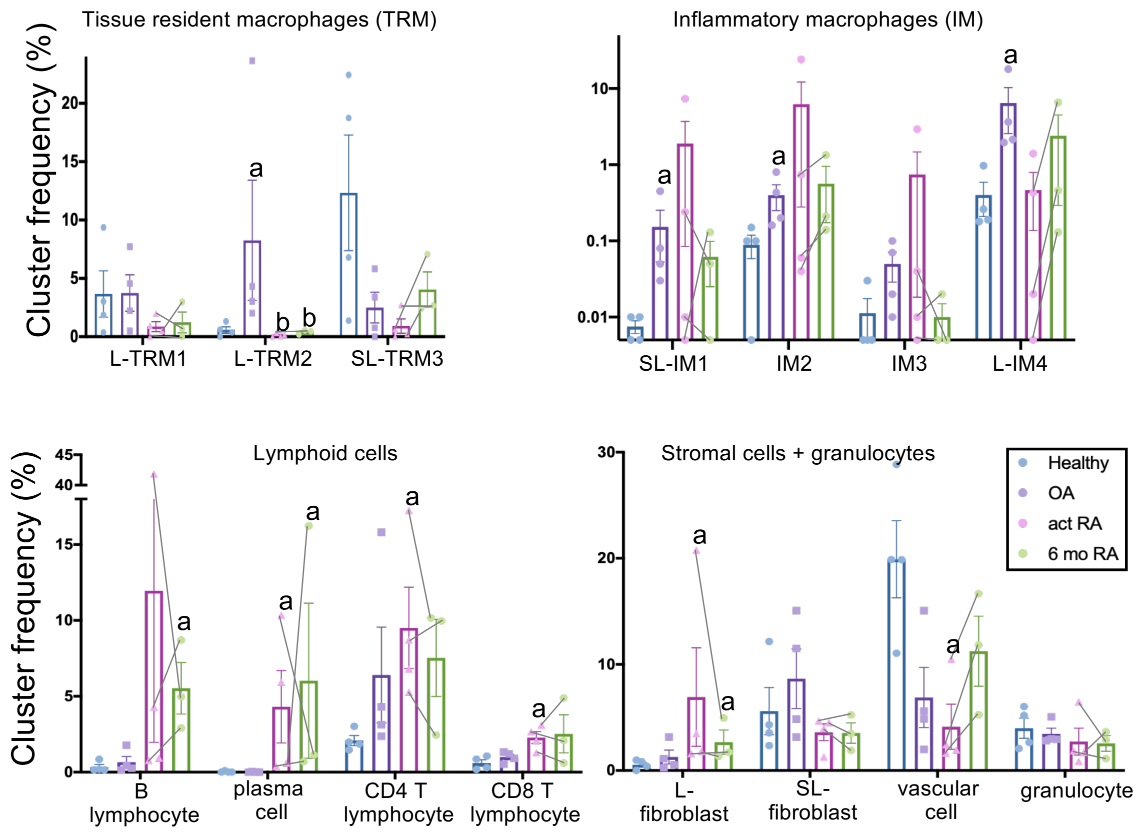


**Fig. S3. Synovial cell frequencies.**

Cell frequency of tissue resident macrophages (TRM), inflammatory macrophages, lymphoid cells, stromal cells and granulocytes is presented in the 4 patients' groups. Mann Whitney tests: a, $p<0.05$ compared to healthy; b, $p<0.05$ compared to OA.

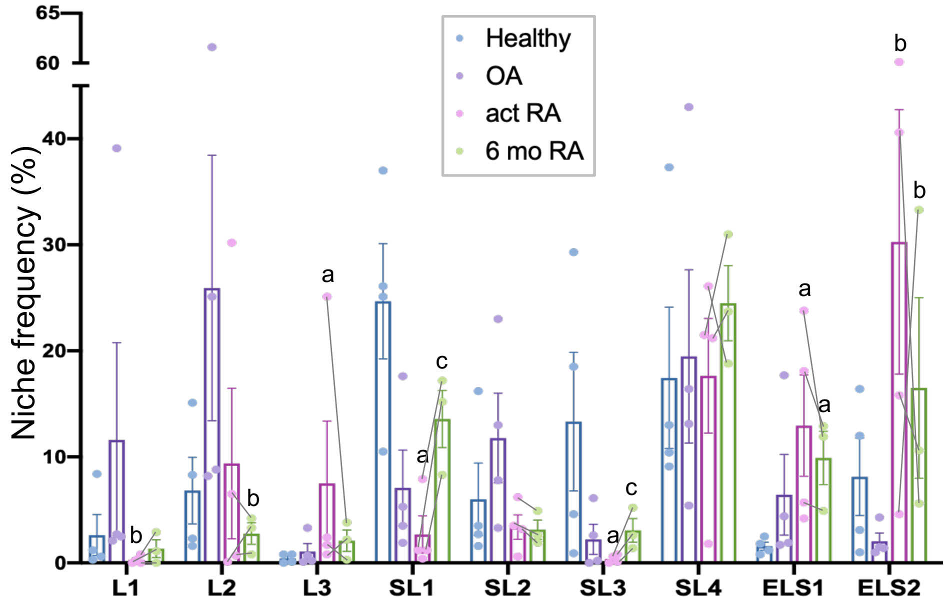


**Fig. S4. Niche frequencies.**
Frequency per sample of all niches in the 4 patients' groups. Mann Whitney tests: a, $p<0.05$ compared to healthy; b, $p<0.05$ compared to OA; c, $p<0.05$ compared to active RA.

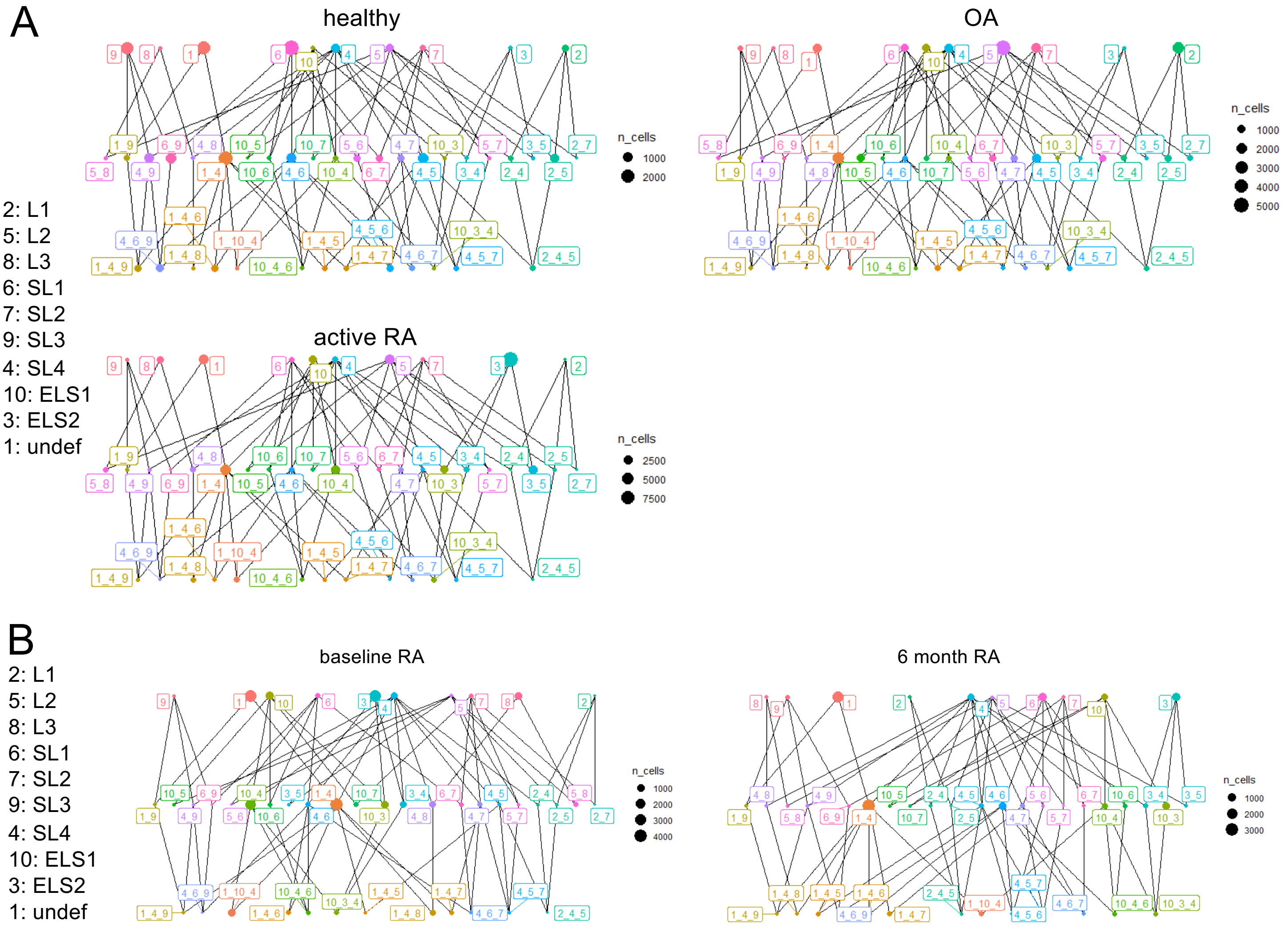


**Fig. S5. Niche interactions.**
**(A)** Interactions of the 10 niches (spatial context) in the healthy (n=4), OA (n=4) and active RA (n=4) patients' groups. **(B)** Interactions of the 10 niches (spatial context) in the active RA at baseline (n=3) and 6 months post-csDMARD treatment (n=3) groups. Dot size represents the number of interacting cells.

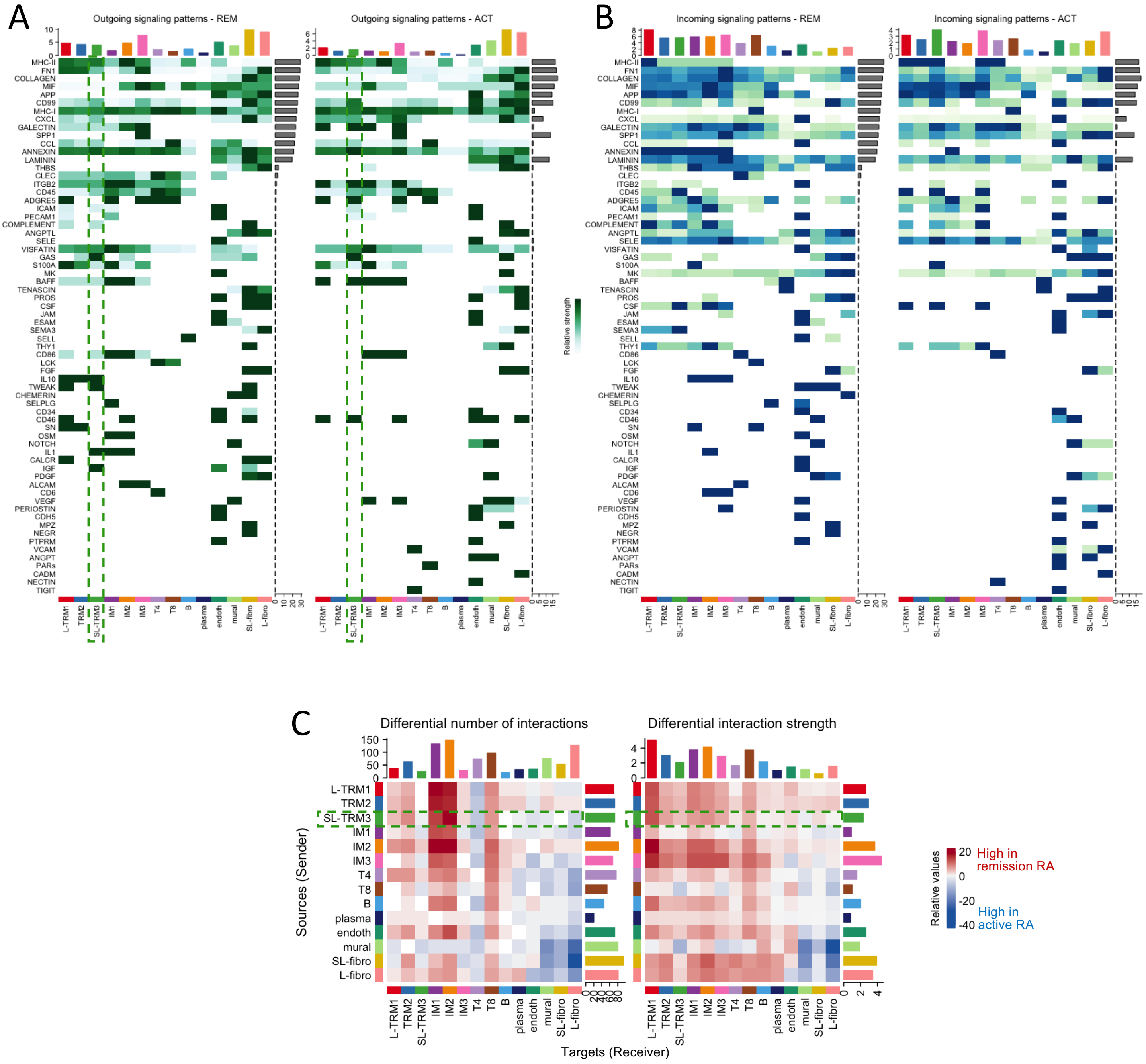


**Fig. S6. Ligand-receptor interactions in active RA versus remission RA synovial cells.**
Heatmaps showing side by side comparison of all outgoing **(A)** and incoming **(B)** signaling patterns in active RA (ACT, n=8) versus remission RA (REM, n=3) synovial cell clusters. **(C)** Heatmap of the differential interactions (number and strength) between synovial cells dissociated from active RA and remission RA tissues. Interactions that are high in active RA are shown in blue, that are high in remission RA in red. Outgoing signaling from SL-TRM3 cells (sources) is highlighted in green.
